# Synthesis and Characterization of ICG-based Near-infrared Photoacoustic Contrast Agents

**DOI:** 10.1101/2025.10.18.683226

**Authors:** Marzieh Hanafi, Giovanni Giammanco, Shrishti Singh, Elsa Ronzier, Aayushi Laliwala, Dana Wegierak, Eric C. Abenojar, Pinunta Nittayacharn, Agata Exner, Parag V. Chitnis, Remi Veneziano

**Affiliations:** Department of Bioengineering, College of Engineering and Computing, George Mason University, Fairfax, VA, 22030; Department of Biomedical Engineering, Case Western Reserve University, Cleveland, OH, 44106; Institute for Advanced Biomedical Research, George Mason University, Manassas, VA 20110; Biomedical Research Laboratory, Institute for Biohealth Innovation, George Mason University, Manassas, VA 20110; Center for Advancing Systems Science and Bioengineering Innovation, George Mason University, Fairfax, VA, 22030; Department of Radiology, Case Western Reserve University School of Medicine, Cleveland, OH, 44106

**Keywords:** Indocyanine green, Photoacoustic imaging, Contrast agent, Near-infrared, DNA nanotechnology, J-aggregates, Lipid nanobubbles, Multi-modal imaging

## Abstract

Near-infrared photoacoustic imaging (NIR-PAI) integrates optical excitation with ultrasound detection to enable high-resolution, deep-tissue imaging by taking advantage of reduced light scattering and absorption in this spectral window. Despite its potential, clinical translation of contrast-enhanced NIR-PAI is limited by the scarcity of effective contrast agents. Indocyanine green (ICG), an FDA-approved NIR dye, is a strong candidate due to its biocompatibility and photoacoustic efficiency. However, its concentration-dependent aggregation, lack of facile targeting strategies, instability in aqueous environments, and low photostability result in variable signal, high background noise, and reduced reliability in vivo. To address these challenges, we developed three biocompatible ICG-based nanoprobe platforms amenable to facile, scalable synthesis: 5-arm DNA-ICG nanostructures (5-arm DNA-ICG), lipid-shelled ICG nanobubbles (ICG-NBs), and Azide-modified ICG J-aggregates (JAAZ). These platforms are designed to preserve ICG monomers or control aggregation, enabling enhanced NIR-PAI performance. Spectroscopic and photoacoustic analyses revealed consistent absorbance and photoacoustic **profiles**, showing enhanced signals compared to free ICG. The greatest improvement was observed for JAAZ, followed by ICG-NBs and 5-arm DNA-ICG. Photostability studies showed that JAAZ aggregation protects ICG from light-induced photodegradation, whereas monomer preservation in 5-arm DNA-ICG and ICG-NBs provides less protection and moderate signal stability. All three probes demonstrated stable performance under physiological conditions, achieved strong signal-to-noise ratios at depth and under tissue-mimicking conditions, and required markedly reduced probe concentrations to generate robust signals. Their modular architectures allow incorporation of targeting ligands, offering molecular specificity and multimodal functionality. Collectively, these contrast agent platforms provide noninvasive, deep-tissue molecular imaging and biosensing, with strong potential for future preclinical and clinical translation, and represent a promising alternative to free ICG for biomedical applications.

## Introduction

Photoacoustic imaging (PAI) is a non-invasive hybrid imaging modality that combines optical excitation with ultrasound detection to enable deep tissue imaging with high contrast and spatial resolution (1–3). In this technique, pulsed laser light excites tissue chromophores, triggering localized heating and thermoelastic expansion. This process generates ultrasonic waves that scatter significantly less than light in biological tissues and that can be detected by a transducer to reconstruct clinically useful images (4–6). PAI is sensitive to endogenous chromophores such as hemoglobin and melanin (7,8) and it has notably enabled visualization of vasculature (9,10) and monitoring of physiological parameters like blood oxygenation, oxygen saturation (sO₂), and oxygen consumption (11). Exogenous contrast agents (CAs) (12) (e.g., organic dyes, gold nanorods, and single-walled carbon nanotubes) (13) can further enhance PAI by enabling targeted imaging with significantly improved contrast. This approach has already been successfully applied to identify tumor lesions (14) and detect lymph node metastases (15).

Imaging depth in PAI can be improved by using near-infrared (NIR) illumination (700 to 1600 nm), as light in this spectral window undergoes reduced scattering and absorption in biological tissues (16). However, widespread adoption of contrast-enhanced NIR-PAI is hindered by limited availability of effective NIR CAs. Indeed, many existing CAs suffer from thermal instability (e.g., gold nanorods (17)), short circulation times (e.g., microbubbles (18)), insufficient biocompatibility (e.g., quantum dots (19)), and non-specific targeting and photobleaching (e.g., small organic dyes (20)), which compromise their imaging performance, especially for *in vivo* applications (21). Indocyanine green (ICG), an FDA approved dye, is a promising candidate due to its NIR absorption peak at 780 nm, biocompatibility, and strong PA signal (22). The utility of ICG for NIR-PAI has already been demonstrated in many clinical settings including for angiography (23), hepatic function assessment (24), and intraoperative imaging (25). Despite its clinical utility and biocompatibility, ICG suffers from low photostability, non-specific tissue accumulation, and limited targeting ability, which results in high background signal (26,27). Moreover, ICG is unstable in aqueous environments and readily forms dimers, trimers, and higher-order structures, particularly at high concentrations (28,29), which leads to variable and unpredictable optical absorption profiles complicating its use in quantitative imaging *in vivo* (30). To overcome these challenges, strategies such as sequestering ICG in nanocarriers have shown promise in enhancing stability, preventing aggregation, and enabling molecular targeting (31).

In this work, we examined three unique strategies to produce ICG-based NIR-PAI contrast agents, which have the potential to provide enhanced stability, reproducible and predictable optical-absorption signatures in biological environments and *in vivo*, and facile conjugation with targeting moieties: 1) tethering ICG molecules on a 2D DNA-based nano-scaffold (DNA-ICG), 2) encapsulation of ICG in lipid nanobubbles (ICG-NBs), and 3) ICG J-aggregates (IJAs).

ICG molecules can be maintained in their monomeric form by templating them with nanoscale precision onto biocompatible DNA-based nanoparticles (DNA-NP) that can be easily designed and self-assembled in 1-, 2-, or 3-D. In addition, these programmable DNA-NPs allow functionalization with various targeting moieties (32,33) to enable cell-specific labeling and enhance molecular imaging applications (34). We previously demonstrated that a specific DNA-ICG structure can be used for voltage-sensing on cell membrane with NIR fluorescence (35). However, the performance of a DNA-ICG contrast agent has not been explored for photoacoustic imaging modality.

Another approach for isolating the ICG molecules is integration of the dye into lipid-shelled gas-core nanobubbles (NBs) (36). Nanobubbles, submicron particles, or ultrafine bubbles (200 - 500 nm) are being explored for photoacoustic imaging, building on their well-established use in diagnostic ultrasound (37). Their lipid shells can be functionalized with ligands to target specific biomarkers and improve accumulation in tissues (38). By incorporating optically absorbing dyes into the shell of nanobubbles, they have the potential to support advanced diagnostic and theranostic strategies through dual-modality imaging (39). In this study, we employed an engineered bubble shell structure that has been shown to provide high stability during repeated and prolonged oscillations, as well as significant variations in shear and turbulence while circulating through the body. This nanobubble-shell design adopts a two-layer architecture with tunable elastic properties, resembling bacterial cell envelopes (40) and filled with Octafluoropropane (C3F8) hydrophobic gas.

In our third strategy we leverage the properties of ICG dyes to form J-aggregates. In J-aggregates, monomeric dyes are arranged in a head-tail configuration and held together by non-covalent bonds (41). Strategies to ensure stability of ICG J-aggregates in circulation and to facilitate functionalization through conjugation typically involve use of nanocarriers (42,43). However, nanocarriers or encapsulation of ICG J-aggregates may restrict tunability in size, targeting, and photoacoustic performance and often require complex synthesis process (44). Here, we exploit a published synthesis method that combines ICG and ICG-Azide dyes (Azide-modified ICG J-aggregates (JAAZ)) to produce monodisperse J-aggregates with direct functionalization capabilities through Azide functional groups.(45) With this synthesis method, careful manipulation of synthesis conditions such as ICG-Azide-to-ICG molar ratio, salt concentration, incubation time and temperature enable precise tuning of the size of the resulting aggregates formed in the range of 230 nm to 1 µm. These particles were also reported to be stable in pH range of 5 to 9 and in 10% FBS solution over time as well (45).

In this study, we present detailed characterization of these three ICG-based CA platforms (Fig. 1) and compare their distinct optical and PA signatures. Their optical stability, modularity, and multimodal imaging capabilities make them promising candidates for noninvasive, deep-tissue molecular imaging and biosensing, with strong potential for future preclinical and clinical translation.

**Figure 1.**
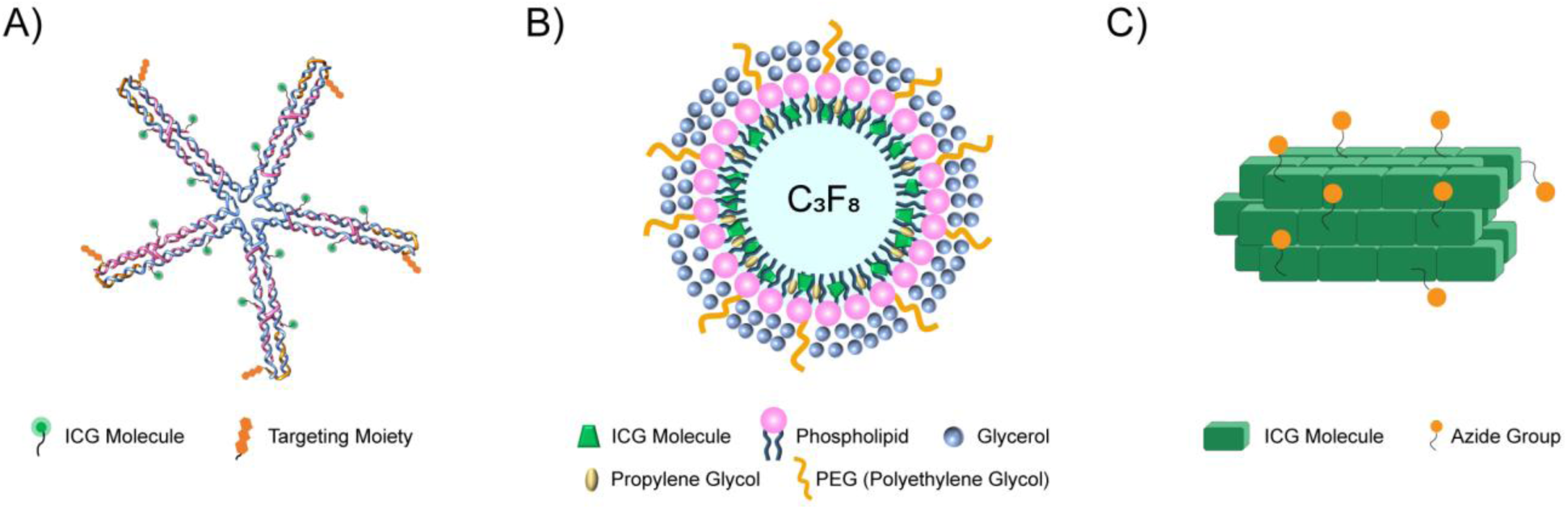
Schematics of the three ICG-based PA probes developed in this study. A) Design of the 5-arm DNA-ICG nanoprobe displaying multiple ICG molecules (green dots) and targeting moieties (orange tri-hexagonal object) (rendering made in Chimera (46)). B) ICG-loaded nanobubbles: The light blue core represents perfluorocarbon gas filled inside, and the green trapezoids display ICG molecules in the lipid shell layer. C) Azide-modified ICG J-aggregates (JAAZ): Each brick represents an ICG molecule. The orange circles indicate Azide functional groups which can be utilized for further conjugation with targeting ligands.

## Results & Discussion

### Synthesis of the ICG-based Contrast Agent Platforms

#### 1. Design and Synthesis of ICG-Templated DNA Scaffolds

DNA nanotechnology offers structural flexibility and allows for creating nanopatterns of ICG molecules on the nanoparticles with control over the inter dye distance (reducing the dimerization risk) and the density of dyes (Figure S1). Building upon our previously published work (35), that used a DX-tile based 1D DNA rod with 4 ICG dyes and two targeting moieties (DIVIN) displayed on the same face, we have redesigned multiple 2D nanoparticles in form of multi-arm structures. These structures have been designed to allow conjugation of 1 to 3 ICG dyes per arm for a maximum number of dyes of 9, 12 and 15 on the 3-arm, 4-arm, and 5-arm, respectively (Figure 1A and S2). The structure used in this study is the 5-arm nanostructure which allowed us to tether the highest number of ICG to maximize the signal intensity. In addition, this new design offers 5 potential sites (one on each arm) for the bioconjugation of the targeting moieties with all the conjugation sites on the nanostructures facing the same direction.

The design for the 5-arm structure is based on a 5-way junction (5WJ), and each structure arm is 73 bp long (or ∼ 25 nm for each arm), following the same design and using an identical set of oligonucleotides to simplify synthesis and reduce the overall cost. The oligonucleotides intended to bear ICG molecules were functionalized at their 3′-ends with Thiol groups and the ICG-modified strands were verified through fluorescence spectroscopy. The inter-dye distance is around 5.5 nm for the three ICG dyes on each arm.

The correct assembly of the nanoparticles and their monodispersity was confirmed via Agarose Gel Electrophoresis (AGE) to verify proper folding and ensure the absence of byproducts through showing a single band on the gel with reference to a 1 kb+ ladder (Figure 2A). Dynamic Light Scattering (DLS) size measurements also confirmed a hydrodynamic size of about 38 nm for 5-arm nanostructures which appears congruent with the values estimated by the structure design (Figure 2D).

**Figure 2.**
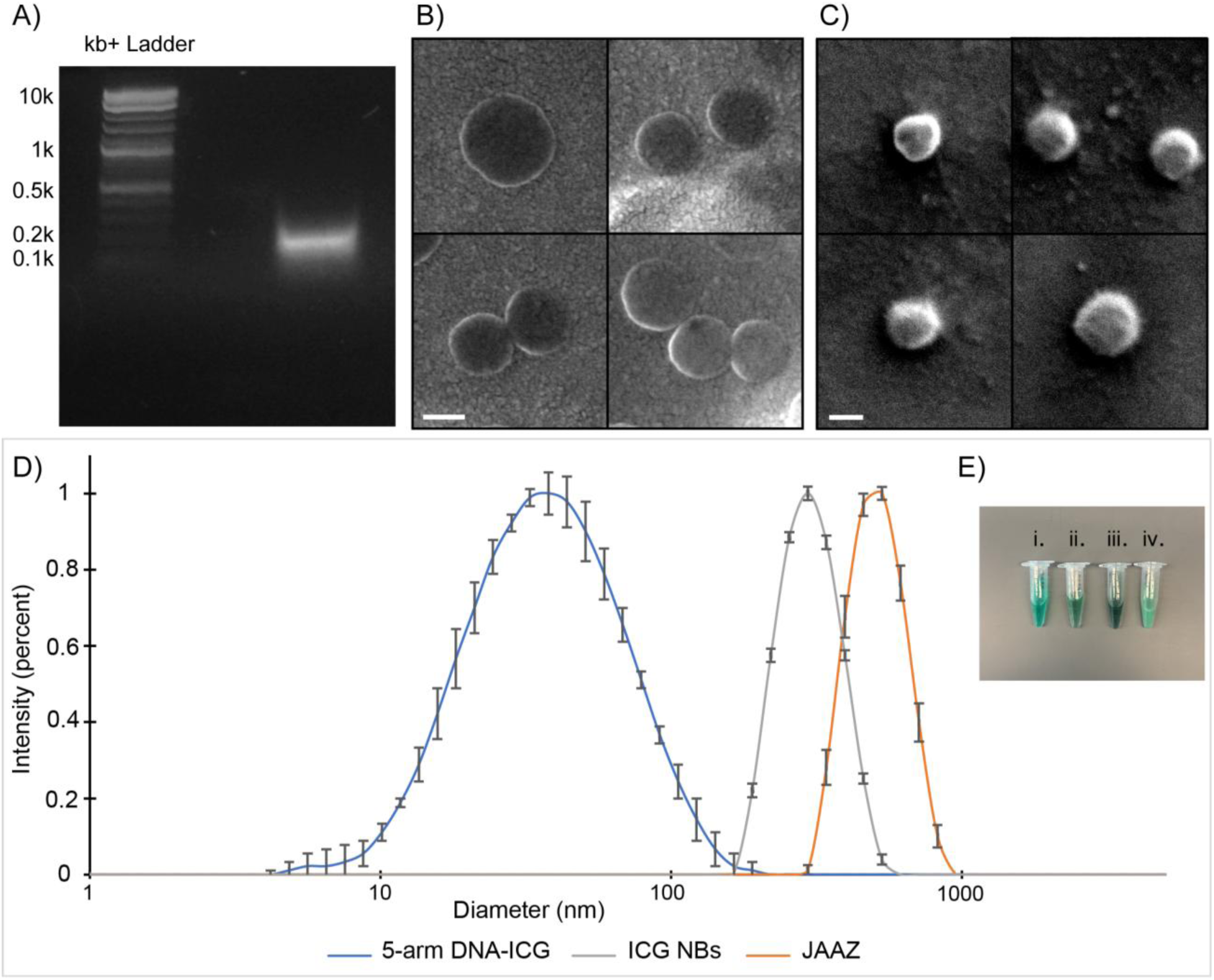
Characterization results for the three ICG-based platforms. A) Agarose gel electrophoresis image for 5-am DNA-ICG probe at 1 µM confirming folding of the correct structure with no byproduct. B) Representative SEM images of ICG NB probe: The dark circles represent the nanobubbles immobilized on the lipid surface. Scale bar: 150 nm. C) Representative SEM images of the JAAZ probes: Scale bar: 400 nm. D) DLS size measurement results depicting a normal distribution centered around 37.84 ± 0.5 nm, 295.3 ± 0.4 nm and 495.49 ± 0.56 nm for 5-am DNA-ICG, ICG NBs and JAAZ respectively. Standard deviation shown as error bars (n = 3 measurements). The intensity values have been normalized to the peak intensity number for each probe. E) Exemplary image of the tubes: i. ICG, ii. 5-arm DNA-ICG, iii. JAAZ and iv. ICG nanobubbles all at 100 µM free dye concentration.

#### 2. Synthesis of Nanobubbles with ICG-Integrated Lipid Shells

To produce the shell material of ICG-loaded nanobubbles, a phospholipid cocktail was prepared with direct incorporation of ICG prior to thermal processing to ensure thorough integration of the dye into the lipid matrix. Following dye-lipid integration, the solution was combined with propylene glycol (serving as an edge activator), glycerol (a membrane stiffener), and phosphate-buffered saline (PBS) to establish a compliant phospholipid layer suitable for bubble formation (47–49). Nanobubble activation was achieved through the injection of Octafluoropropane (C₃F₈), followed by mechanical agitation.

DLS size analysis of the ICG-incorporated NBs demonstrated a normal distribution centered at ∼295 nm in diameter, proving the bubble sizes to be within the expected nanometer range, indicating successful control over bubble formation. (Figure 2D). The concentration and buoyancy of the bubbles tested using resonant mass measurement (RMM) showed average 4.15 x 10^11^ bubbles/mL that comprised of 92.4% positively buoyant particles, thus confirming the formation of stable nanobubbles (Figure S3). SEM imaging further validated the morphology, showing spherical structures with smooth surfaces, which is consistent with typical phospholipid-based nanobubbles and confirms the size measurements further as well (Figure 2B and S4a).

#### 3. Controlled Aggregation of ICG Molecules and JAAZ Synthesis

We exploited the synthesis method for JAAZ production presented in Singh et al.’s work (45) in which they showcase monodisperse, nanometer-sized, functionalizable Azide-modified ICG J-aggregates (JAAZ) acquired through a facile synthesis method without the further need for syringe filtration or encapsulation in nanocarriers. They took advantage of the presence of KCl salt since monovalent metal cations are reported to promote the formation of J-aggregates in anionic cyanine dyes like ICG by reducing electrostatic repulsion among them (50,51).

As mentioned before, the size of the resulting JAAZ can be modulated by fine-tunning experimental factors. For this study, we aimed for using ∼500 nm JAAZ particles obtained with the following synthesis parameters: 1:10 ICG-Azide:ICG molar ratio, total dye concentration of 1 mM, 20 h of incubation at 60 °C and 1 mM of KCl.

DLS and SEM were performed on JAAZ to assess the size distribution and acquire visual confirmation of particle morphology. In SEM images JAAZ particles appear as small circular nanoparticles with an average diameter of 468.8 nm (Figure 2C and S4b). DLS measurements demonstrated a normal distribution of JAAZ particles diameter centered at ∼495.5 nm (Figure 2D), indicating consistent particle formation and suggesting good control over synthesis parameters. Together, these results confirm that the JAAZ particles exhibit both size uniformity and morphological consistency, which are essential for their further applications.

### Optical Characterization of the Probes

#### 1. Optical Absorbance Measurements

The absorbance spectra of all three newly developed ICG-based contrast platforms were examined in PBS with absorbance spectroscopy for the wavelength range of 500-900 nm. ICG’s absorbance was measured through the same procedure as the reference to compare the probes and observe the change in their absorbance readings relatively. All measurements were carried out for five different free ICG concentrations of 15, 30, 60, 90 and 120 µM.

As shown in Figure 3A, the plain ICG shows two absorbance peaks in its spectrum at 710 and 780 nm representing its dimeric and monomeric form (52) with the latter possessing higher absorption wavelength (53). Even for concentrations as low as 30 µM the dimeric peak seems prominent and only for the concentration of 15 µM, the monomeric form appears to exist in higher quantities in the solution. This behavior of ICG had been shown before in several literature demonstrating the concentration dependent aggregation of ICG molecules in relatively high concentrations (28). The peak for the dimeric form of ICG keeps getting more prominent with increasing the concentration of ICG. Moreover, sample absorbance rises with ICG concentration with the highest optical density at about 1.5 at dimeric peak and about 1 for the monomeric peak for the highest ICG concentration (120 µM).

**Figure 3.**
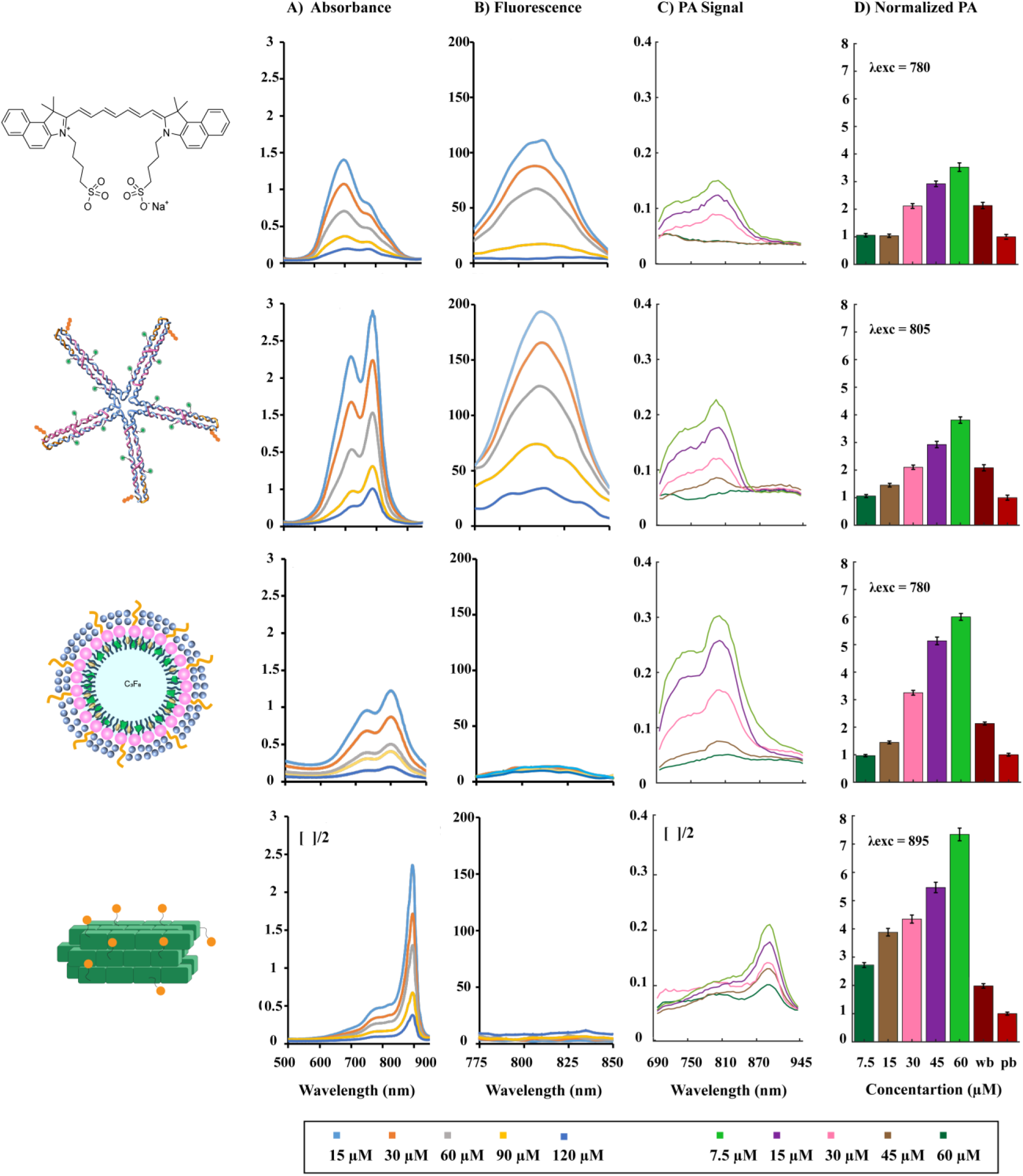
Optical properties (at concentrations 15, 30 60, 90 and 120 µM) and photoacoustic signal measurements (samples mixed with blood at a 1:1 ratio; final concentrations 7.5, 15, 30, 45, 60 µM) of ICG-based probes, namely 5-arm DNA-ICG, ICG nanobubbles and JAAZ compared to free ICG dye. A) Absorbance profile across wavelengths from 500 to 900 nm. B) Fluorescence intensity measurement conducted over a wavelength range of 775 to 850 nm. C) Photoacoustic spectra at multiple wavelengths spanning 690 to 945 nm. D) PA amplitude at each CAs peak absorbance normalized to PA amplitude of a 1:1 PBS/blood sample.

Moving forward to the absorbance readings for 5-arm, the graph denotes that conjugating DNA nanoparticles with ICG resulted in slight red shift in the absorption peak (monomeric form) to 805 nm at all concentrations which lies in a biologically favorable window (near infrared) due to minimal background absorption from endogenous chromophores. A drastic decrease in the dimer component is observed, which indicates that ICG remained in monomeric form mostly, even at high concentrations where dimerization and aggregation is observed for free ICG. This phenomenon happened as a reason of controlling inter-dye distance through the DNA scaffolded architecture to avoid dye-to-dye interactions that lead to dimerization, a blue peak absorbance shift and inconsistent optical performance (54). In addition, the optical density for the prominent peak which belongs to the monomeric form rose to about 3 for the highest concentration of ICG which is doubled compared to free ICG at its prominent peak.

ICG-nanobubbles demonstrated an optical absorption signature similar to ICG dye in terms of optical density amplitude and two different peaks for monomeric and dimeric forms of ICG molecules. The non-negligible distinction of the ICG nanobubbles absorption spectra show a higher absorption peak at 780 nm, which belongs to the monomeric form of ICG, at all concentrations without a shift to the 710 nm peak as the prominent one. This observation points out the capability of decreasing the formation of dimeric form of the ICG molecules when incorporated into lipid nanobubbles.

Looking at JAAZ absorption spectra, there is no sign of the peaks for either monomeric or dimeric form of the ICG since we are dealing with ICG aggregates instead of free molecules. The graph depicted a strong, red-shifted absorption peak at 895 nm. The single absorption peak of JAAZ is much narrower compared to that of ICG, indicating an improved signal to noise ratio (SNR). In addition, JAAZ absorption spectra demonstrate a rocketed optical density for all the concentrations with the highest peak absorption of 5 indicating a more than 3-fold increase compared to ICG (∼1.5) for the equivalent ICG concentration of 120 µM. This is a result of an increased dye local density due to the formation of J-aggregates. The absorbance comparison of the probes demonstrates that structural modification of ICG significantly improves its optical properties, exhibiting stronger and more consistent signals.

#### 2. Fluorescence-Photoacoustic Bimodality of the Probes

Fluorescence spectrometry revealed key differences among the three ICG-based contrast agents when compared to free ICG. The measurements were conducted for the concentrations same as optical absorbance measurements for the wavelength region of ∼775–850 nm via a fluorescence plate reader and were compared to plain ICG. Figure 3B indicates an increase in fluorescence intensity as a function of ICG concentration in 5-arm DNA nano-constructs compared to ICG fluorescence spectra. The fluorescence intensity of DNA nanoprobes were improved by two-fold at the peak for the samples with equivalent ICG concentration of 120 µM. This probably happened due to the change in the electrical charge of the probe since DNA scaffolds bear negative charges (55). This also makes DNA nanoprobes multi-modal contrast agents that can be utilized for both NIR fluorescence imaging and photoacoustic imaging.

ICG-nanobubbles exhibited diminished fluorescence signal amplitudes, likely due to dye incorporation within the lipid shell. However, they can still present ultrasound imaging results as contrast agents designed for measurements performed through this imaging modality. Similarly, JAAZ showed almost no fluorescence upon optical excitation, suggesting strong fluorescent quenching effects, which are conducive to PA-based sensing and imaging where reduced fluorescence improves PA signal efficiency.

### Photoacoustic Signal Intensity Results

#### 1. Photoacoustic Signal Amplitude Measurements in Blood

To simulate intravenous delivery, the probes were characterized in blood by mixing same proportions of probe solutions in PBS and whole sheep’s blood. The PA signals for the four contrast agents were recorded at 690–945 nm wavelengths and their signal intensity were measured at their absorption peak wavelength at resulting concentrations of 7.5, 15, 30, 45, and 60 µM. These measurements were carried out via our developed photoacoustic experiment setup in the laboratory (Figure 5).

In order to collect the PA data, the samples were inserted in tubing and submerged in deionized (DI) water. To maintain a frequency comparable to that of transducers used for photoacoustic imaging in small animals, a focused piezoelectric transducer operating at 35 MHz was positioned above the sample at its focal distance and with its functional element immersed in the water. The samples were excited at an oblique angle using a wavelength-tunable pulsed laser.

Our results as shown in Figure 3C confirmed that the photoacoustic spectra of these contrast agents agree with the absorbance measurements which firstly means preserved peak absorbance wavelengths and secondly higher PA amplitude for all the probes compared to ICG for peak absorbance wavelengths with highest being for JAAZ followed closely by ICG nanobubbles and then DNA ICG nanostructures. Concentration-dependence of PA signal of all probes was also conserved when nano-probes were introduced into the blood.

PA signal intensity of all probes at their absorption peak wavelength normalized to the PA of 1:1 mixture of PBS/blood for the mentioned concentrations are shown in figure 3D. All three nanoprobes generated higher PA signals than whole blood at concentrations ≥30 µM. The most pronounced gains in all developed probes’ PA signal amplitudes occurred at relatively high concentrations (>90 µM). Among them, JAAZ produced the strongest signal, exceeding that of whole blood even at 7.5 µM, and showing more than a fivefold increase at 60 µM as opposed to ICG. This is attributed to its high local dye density and red-shifted absorption peak, which enhance PA sensitivity and spectral distinction. ICG-nanobubbles showed a fourfold increase in PA signal at 60 µM, with enhanced signal in blood in contrast to their signal while dispersed in PBS only (Figure S5). This outcome suggests that lipid encapsulation stabilizes the monomeric form of ICG and improves acoustic response, potentially due to either direct interactions or hitchhiking on red blood cells (56). The DNA-ICG constructs preserved a linear relationship between concentration and PA signal, with consistent monomeric absorption retained at all tested concentrations. This stability results from the spatial control over dye placement afforded by the DNA scaffold, effectively preventing dye dimerization. In contrast, free ICG demonstrated limited PA signal enhancement beyond 60 µM consistent with its tendency to form dimers and lacked a detectable peak at lower concentrations. Overall, the results highlight the improved PA performance and in-blood stability of the engineered nanoprobes, with JAAZ offering the strongest signal, followed by nanobubbles and DNA-based constructs.

#### 2. Photoacoustic Signal Data for Different Probe Concentrations

All previously reported concentrations of the probe samples referred to the equivalent free ICG dye content, and despite identical ICG concentrations, all developed probes demonstrated enhanced photoacoustic signal intensity. However, one of the greatest achievements in developing the ICG-based platforms covered in this research was decreasing “probe concentrations” remarkably while improving the PA signal amplitude compared to plain ICG dye and still providing adequate visualization for most clinical applications. Decreasing the CA concentration and lowering the number of injected particles in intravenous injections has been proven to be beneficial in terms of decreased risk of complications and potential adverse effect of IV administration especially for high-risk patient groups (57,58). As discussed before, every 5-arm design allows the particle to carry up to 15 ICG dye molecules, effectively concentrating signal within fewer particles. JAAZ and nanobubble particle concentrations were quantified using ZetaView^®^ and resonant mass measurement (RMM), respectively.

In figure 4A, the y-axis represents the photoacoustic (PA) amplitude of all probes at equal (60 µM) dye concentration, normalized to that of the free ICG probe. The x-axis shows the calculated probe concentrations for each CA formulation based on their particle molar concentrations. The data reveals a significant reduction in required probe concentrations for the developed contrast agents, particularly for JAAZ followed by ICG NBs, attributed to their larger structures. Despite lower particle doses, these agents achieved superior signal intensity, underscoring their potential as safe, high-performance candidates for NIR photoacoustic imaging.

**Figure 4.**
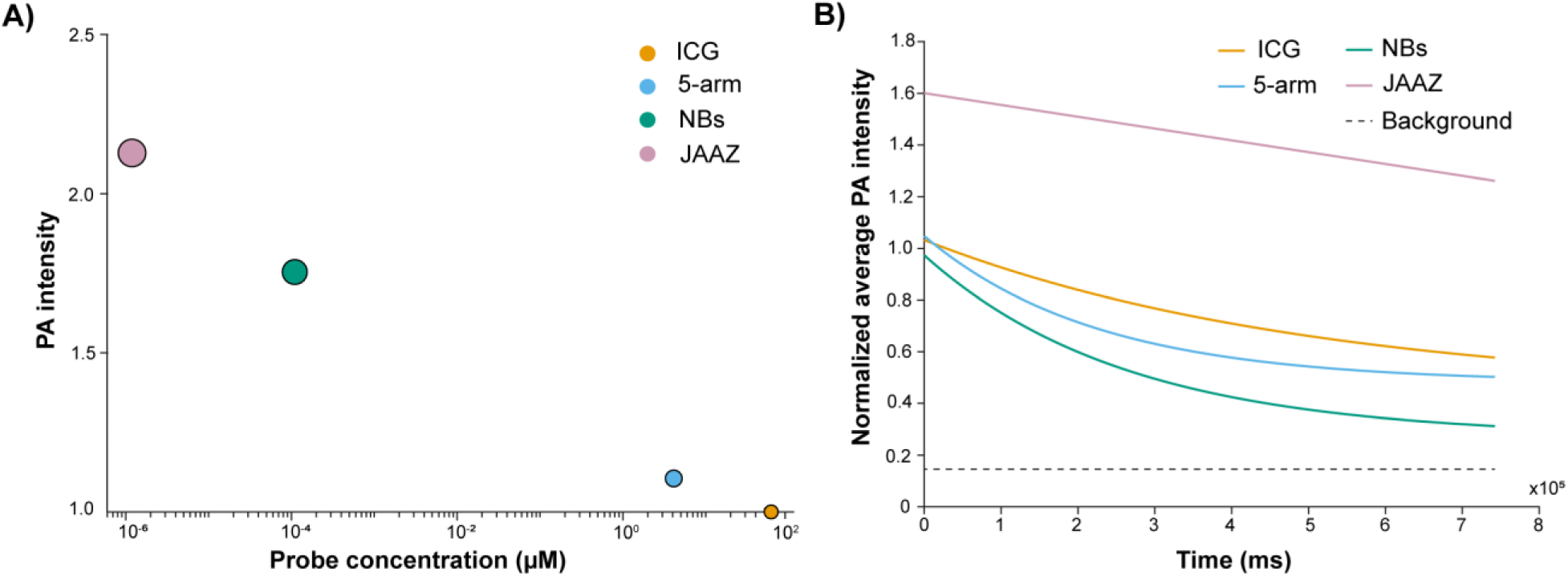
PA Response and Temporal Decay of ICG Platforms A) Logarithmic plot of the PA intensity of ICG, 5-arm DNA-ICG, JAAZ and ICG nanobubbles at an equivalent dye concentration of 60 µM normalized to ICG PA amplitude vs. probe molar concentration of the samples in µM. Marker sizes are proportional to probe particle sizes on a logarithmic scale for visual emphasis. B) PA signal magnitude of the developed platforms over the period of about 12 minutes, normalized to ICG PA signal intensity at time zero (t_o_) and fit to an exponential decay function. All agents were in PBS at an equivalent ICG concentration of 50 µM. The dashed black trend line represents the signal measured in the background.

**Figure 5.**
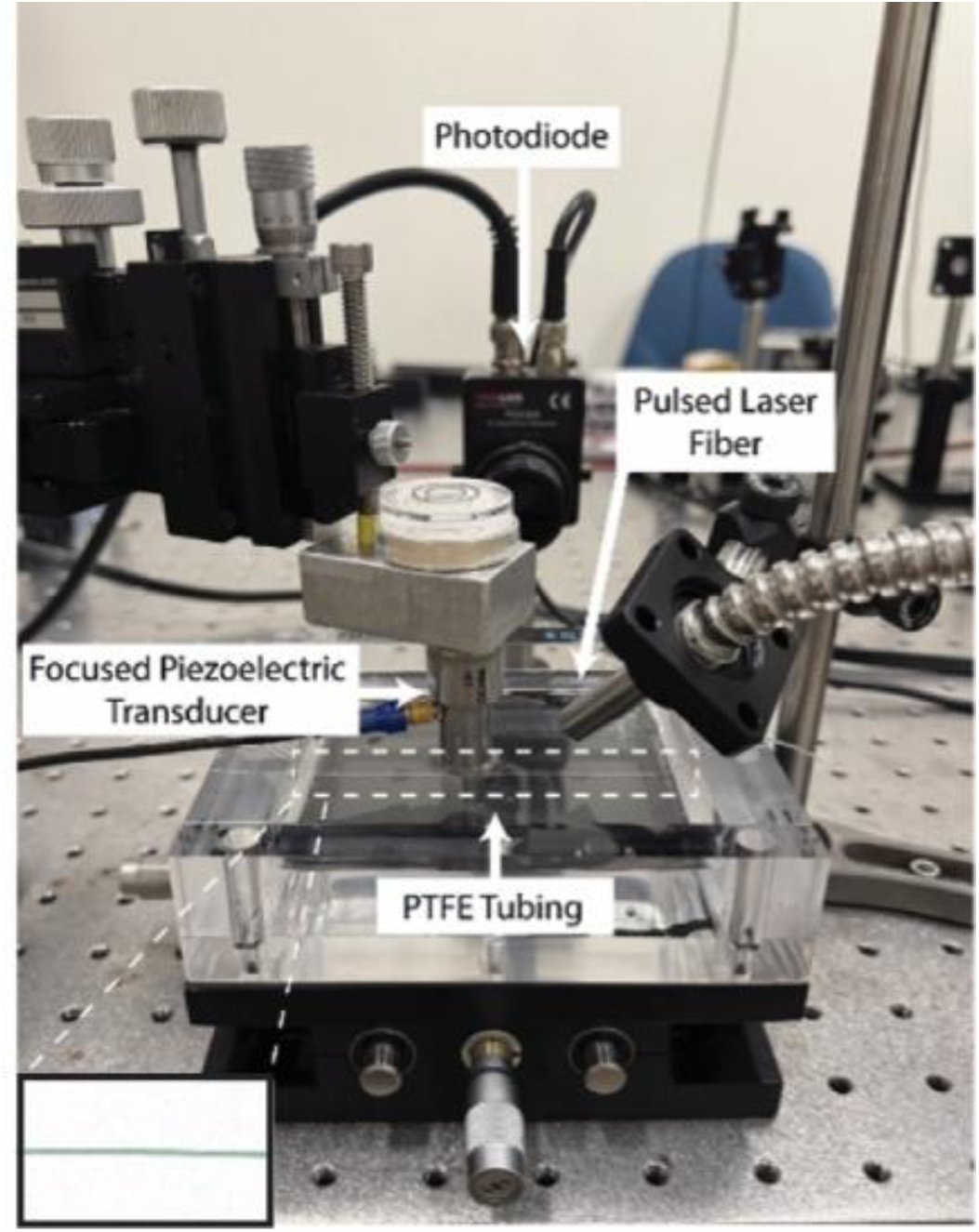
Custom photoacoustic setup developed by the lab. The inset image demonstrates a PTFE phantom tube filled with a sample probe.

#### 3. Photostability and SNR Evaluation Under Tissue-Mimicking Conditions

To further assess the synthesized CAs and determine the stability of the probes upon laser irradiation, we conducted a timeseries stability assay of the probes using PA imaging. Utilizing Fujifilm VisualSonics Vevo F2 device, the PA signal intensity of all probes at comparable concentrations of 50 µM in PBS was measured over 2,970 frames equal to 742 seconds (∼12 minutes). Accordingly, to evaluate the performance of the modified ICG-based contrast agents in *in vivo* conditions the samples were introduced in tubing, stabilized and submerged in a chamber filled with DI (deionized) water and an organic chicken breast tissue layer of 3.5 mm thickness was placed in between the tubing and a 29 MHz transducer (∼1 cm overall distance) to provide tissue-mimicking conditions. They were subjected to 20 laser pulses in a second (20 Hz) at their peak absorbance wavelength which leads to a total of 14,840 laser pulses for the experiment time. The PA overtime for all the probes was measured and averaged along the length of the tube and the resulting data were modeled using a single exponential decay model with an offset. The decay constant (τ), as a measure to verify the pace of signal decay, was calculated for all the probes based on the fit model (Table 1). Background was defined as separate ROI (region of interest) for each contrast agent measurement, placed in the surrounding chicken tissue. Figure 4B depicts a decreasing trend for all the probes as expected compared to the background while DNA-ICG and ICG nanobubbles present a slightly steeper downward trend in comparison to free ICG dye. As JAAZ shows increased initial PA signal magnitude, it also highlights a much milder and linear decline in signal intensity, only losing 22% of the initial signal after 12 minutes of constant irradiation. This outcome was confirmed by calculation of the decay constant (τ) (Table 1), revealing that while JAAZ possesses a distinctly pronounced photostability, 5-arm and ICG-NBs are more prone to photobleaching relative to ICG’s performance under PAI (59). The underlying reason for the photobleaching outcome in general could be the presence or absence of aggregates. Owing to the aggregated structure of JAAZ particles, the ICG molecules are less susceptible to light-induced degradation compared to their individual monomeric state (60,61). In a similar manner, the lower quantity of ICG dimers in 5-arm and ICG-NBs results in less protection of the probe from degradation. This phenomenon further proves the isolation of ICG monomers through nanopatterning the molecules on DNA scaffolds or their integration into lipid nanobubbles (Figure S6 and S7).

**Table 1.** Photostability and Signal-to-Noise Ratio of the Three ICG-based Nanoprobes.

| Contrast Agent | ICG | DNA-ICG | ICG NBs | JAAZ |
| --- | --- | --- | --- | --- |
| Normalized PA Signal Amplitude at $t_0$ | 1 | 1.198 | 1.009 | 1.607 |
| $\tau$ (s) | 506.3 | 224.9 | 262.7 | 252,100 |
| SNR (dB) | 51.27 | 51.43 | 49.50 | 53.53 |

\*The table exhibits the initial PA signal amplitudes normalized to that of free ICG, the calculated decay constants based on the same fit model ( $\tau$ ) and the SNR calculated for the probes inserted into tubing in a water media and under a tissue layer of 3.5 mm.

We investigated the signal-to-noise ratio (SNR) of the developed CAs in the same fashion and calculated the numbers in Decibels (dB) through defining the mean of the signal for the first 100 frames (20 seconds) and the standard deviation of the noise for the same time duration. The background noise was determined as the signal intensity within an ROI on the chicken tissue layer which was used to simulate a biological tissue. All agents showed strong SNR quantities of about or higher than 50 dB while displaying an SNR ≳ 30 dB is considered high quality for contrast-enhanced PA imaging *in vivo* (62–64).

## Conclusion

While indocyanine green (ICG), an FDA-approved NIR dye, represents a promising candidate as a CA for NIR-PAI, its concentration-dependent optical absorption and lack of facile targeting strategies limit its use as an NIR-PA contrast agent. To address this unmet need, we developed and examined novel strategies for synthesizing ICG-based contrast-agent platforms that either preserved the monomeric form of ICG on nanoparticles or produced controlled monodisperse aggregates. The resulting NIR-PA imaging agents were DNA-ICG nanoprobes (5-arm), ICG-nanobubbles (ICG NBs) and ICG-Azide J-aggregates (JAAZ). DNA-ICG platform consisted of biocompatible DNA-based nanoparticles that used a tile assembly method and biorthogonal modifications to precisely template ICG molecules onto programmable DNA scaffolds, enabling structural flexibility and nanoscale control. ICG nanobubbles were lipid-shelled, gas-core nanoparticles filled with Octafluoropropane gas and stabilized by lipid shell, that encapsulates ICG dye molecules. Azide-modified ICG J-aggregates formed by combining ICG and ICG-Azide dyes under high-temperature incubation with KCl resulted in monodisperse particles with tunable size and direct functionalization capabilities.

Our results demonstrate that all contrast agents produced a stronger NIR absorption through higher local dye concentrations and PA signal amplitudes greater than that from whole blood at concentrations above 30 µM and as low as 7.5 µM for JAAZ, enabling deeper tissue imaging. JAAZ, ICG NBs and 5-arm DNA-ICG demonstrated higher PA signal in blood in descending order compared to ICG at similar concentrations in addition to showing structural stability in physiological conditions. Notably, probe concentrations were reduced by orders of magnitude for the new probes, yet still enhancing the PA signal amplitude, offering sufficient visualization for most clinical applications with markedly fewer injected particles.

In comparison to free ICG, the defined nanoscale geometry of the DNA scaffolds allowed us to isolate ICG molecules in distinct positions, enabling predictable absorbance behavior and minimizing dimerization and self-quenching of the ICG molecules. Similar to free ICG dye, DNA-ICG probes are bimodal and can be used for optical fluorescence and PA imaging. JAAZ showed a strong, red-shifted peak absorbance with a profile distinctly different from endogenous chromophores, which can facilitate convenient spectral unmixing and deep-tissue targeted imaging. JAAZ and ICG-nanobubbles exhibited reduced fluorescence signals making them particularly conducive for PA-based sensing and imaging. ICG-nanobubbles are acoustically active thus considered bimodal as well and can be used for both ultrasound and PA imaging. JAAZ unveiled a pronounced photostability whereas DNA-ICG and ICG-NBs demonstrated a modest reduction in photostability relative to ICG which is attributed to the increased ICG monomer content. All contrast agent platforms displayed adequate SNR values at depth and under tissue mimicking conditions.

Since these nanoprobes incorporate FDA-approved ICG, biocompatible DNA, lipids and gas, they are all inherently biocompatible and suitable for clinical translation. The ability to incorporate cell-targeting ligands in all presented CAs can provide high cell specificity, further enhancing the probes’ effectiveness in molecular imaging applications. Moreover, These CA platforms offer key advantages such as facile and scalable synthesis and size-tunability. Each of the designed probes would be an ideal NIR-PA contrast agent variant depending on the end application. While JAAZ indicated a superior performance in PAI, ICG-nanobubbles demonstrated not significantly different PA signal amplitudes in blood and can enable dual modality imaging. Meanwhile, it is only DNA-ICG probe which while maintaining a comparable PA signal to ICG, achieves amplified fluorescence performance, allows for precision nano-engineering, structure organization and multiplexing potentials.

Taken together, these configurations’ structural and functional frameworks mitigate the known challenges of free ICG as a CA while fully leveraging its clinical advantages within modular designs. They hold strong potential to overcome current NIR-PA contrast agent limitations and facilitate the clinical translation of next-generation molecular imaging technologies, representing a promising alternative to free ICG for biomedical applications.

## Materials and Methods

### 1. Materials

ICG and ICG Azide were purchased from AAT Bioquest for the production of JAAZ and ICG-Maleimide was acquired from Diognocine (BioActs). ICG used for ICG NBs synthesis was purchased from MP biomedicals. Dimethyl Sulfide (DMSO) and N,N-Dimethylformamide (DMF) were bought from Fisher Chemical and VWR Life Science respectively. Oligonucleotides and modified oligonucleotides (thiol modified) used for DNA scaffolds assembly were obtained from Integrated DNA Technologies (IDT) and utilized without additional purification. All sequences used are listed in the Supplementary Tables S1. The folding buffer used for assembly of the strands is composed of TAE buffer (40 mM TRIS, 20 mM acetic acid, 2 mM ethylenediaminetetraacetic acid) complemented with 12 mM magnesium chloride (MgCl_2_) at pH 8.0. For gel electrophoresis, high-melt agarose (IBI Scientific), ethidium bromide (Invitrogen) and Gel Loading Dye, Purple (6X) (New England BioLabs) were used. SEM samples were mounted onto a carbon tape (Electron Microscopy Sciences) on aluminum pedestals (RS-MN -10-005112-50; RaveScientific). Afterwards, they were flash-frozen in liquid nitrogen (Roberts Oxygen) and freeze-dried in a Millrock Technology device. Phosphate buffer saline (PBS, pH 7.4) at 1X was used for the dilution of the ICG, DNA strands and all the developed ICG-based probes (Sigma-Aldrich). TCEP HCl (Tris(2-carboxyethyl)phosphine hydrochloride, VWR Life Science), Sodium Acetate (VWR Chemicals, Sterile Solution, pH 5.2), Absolute Ethanol (Fisher Bioreagents) and Isopropyl Alcohol 99.5% (VWR Chemicals) were utilized in the modification process of the oligonucleotides modified with ICG Maleimide. The Amicon Ultra-0.5 Centrifugal Filter MWCO Millipore and Potassium Chloride (KCl) were purchased from Sigma-Aldrich. DBPC (1,2-dibehenoyl-sn-glycero-3-phosphocholine), DPPE (1,2-dipalmitoyl-sn-glycero-3-phosphoethanolamine) and DPPA (1,2-dipalmitoyl-sn-glycero-3-phosphate) were purchased from Avanti Polar Lipids Inc, and mPEG-DSPE2000 (1, 2-distearoyl-sn-glycero-3-phosphoethanolamine-N-[methoxy(polyethylene glycol) 2000] from Laysan Lipids. Propylene glycol (PG) was purchased from Sigma Aldrich. Glycerol (Acros Organics) was purchased from Thermo Fisher Scientific and Octafluoropronane (C3F8) was acquired from Airgas USA LLC. Whole sheep’s blood containing anticoagulant used for PA signal measurements was purchased from Hardy Diagnostics. Sterile nuclease-free water from VWR Life Science was used in any step in the probes’ synthesis process that involved water. The low-volume quartz glass cuvettes for DLS were obtained from Malvern Panalytical.

### 2. Methods

#### 2.1. Synthesis of the Nanoprobes

##### 2.1.1. “5-arm” DNA-ICG Nanoprobes

###### 2.1.1.1. DNA Nanoparticle Design

The DNA nanoparticles were designed using the Tiamat software (65). The sequences of the various strands were carefully designed to ensure orthogonality, aiming to enhance folding efficiency and prevent mispairing. The sequence orthogonality was confirmed using NCBI Blast (66,67).

###### 2.1.1.2. DNA Oligonucleotides Conjugation with ICG

ICG-Maleimide was resuspended in DMSO/DMF and further diluted with PBS 1X while keeping the final concentration of the resuspension solvents below 8%. Selected DNA strands on the DNA nanoparticles (3 per arm) were modified with ICG via thiol-maleimide chemistry. Briefly, DNA strands modified with Thiol groups at their 3’ end (Table S2) were reduced with a 20x excess molar ratio TCEP HCL for 15 minutes and then directly mixed with 10x excess molar ratio of ICG-maleimide without removing TCEP and kept for 2 hours at room temperature (RT) and overnight at 4 °C. Sodium acetate at the final concentration of 0.3M was used for the precipitation of DNA-ICG followed by the addition of isopropanol at 60% (ice cold, 99% purity). The mixture was centrifuged (Thermo Scientific Sorvall ST 8R centrifuge) for 4 hours at 1 °C (30,000 × g). The pellet was washed with ice-cold absolute ethanol and centrifuged under the same setting for an additional 15 minutes. After removing the ethanol, the pellet was dried under the vacuum in the dark and resuspended in water. The efficiency of ICG conjugation was determined via spectroscopic evaluation using absorbance and fluorescence measurements (Tecan Safire 2 Multi-Detection Plate Reader and Molecular Devices Spectramax Gemini Em Fluorescence Microplate Reader).

###### 2.1.1.3. “5-arm” DNA Structure Assembly

The 5-arm DNA nanoparticles were assembled by mixing the modified or non-modified oligonucleotides at the specific molar ratios presented in the Table S1 and S2, in folding buffer and a final DNA nanoparticle concentration of 10 µM. The solution was then slowly annealed from 95 °C to 4 °C over 2 hours in a thermocycler (T100 BioRad) for proper hybridization of the strands. Following folding, the nanostructures were stored at 4 °C without further purification prior to characterization. The concentration of the DNA strands and nano-probes was measured by a Microvolume UV-Visible spectrophotometer (NanoDrop One from Thermo Scientific). The final 5-arm nano structure consists of 766 base pairs with a molecular weight of ∼0.4 MDa.

##### 2.1.2. ICG-Nanobubbles Production

Indocyanine green nanobubbles (ICG NBs) were formulated using a previously reported method (40,68,69). Briefly, phospholipids including DBPC, DPPE, DPPA, mPEG2000-DSPE (6:2:1:1), and ICG (0.3) were dissolved in propylene glycol. After complete dissolution, a solution containing glycerol in PBS was added to the lipids and sonicated to form a lipid emulsion. The lipid emulsion was aliquoted to a 3 mL glass vial that was sealed, and the headspace of the vial was exchanged with Octafluoropropane (C3F8) gas. Subsequently, self-assemble bubbles were formed by shaking the vial for 45 s using Vialmix® and ICG NBs were isolated by differential centrifugation (Eppendorf centrifuge 5804) at 50 G for 5 min. The final product contains ICG at a theoretical concentration of 0.3 mg/mL in the solution. The nanobubbles were characterized for size using dynamic light scattering (DLS), concentration and buoyancy (resonant mass measurement), and scanning electron microscopy (SEM). The vials are protected from multiple freezing-thawing processes and stored at -80 °C prior to further use.

##### 2.1.3. JAAZ Formation

Following the protocol developed by Singh et al. (45), we assembled JAAZ particles using a 1:10 molar ratio of ICG-Azide:ICG in water with 1 mM of KCl for a total dye concentration of 750 μM and incubated the mixture at 60 °C for 20h. The JAAZ particle samples underwent purification with 0.5 mL Amicon centrifugal filters (100 kDa cutoff) to eliminate unaggregated ICG monomers. The filtration process involved 3 steps of centrifugation at 4000 × g for 5 minutes at room temperature to ensure thorough removal of unreacted dye. The samples were stored at 4°C prior to further use and later diluted in PBS.

#### 2.2. Nanoprobes Characterization

##### 2.2.1. Agarose Gel Electrophoresis (AGE)

The folded 5-arm probes were diluted to the concentration of 1 μM and mixed with 1x Gel Loading Dye before being loaded onto 50 mL of a formed high-melt gel at 1.2% in 1x TAE (pH 8.0) and pre-stained with 0.5 µg/mL Ethidium Bromide. A Kb+ ladder was used, and electrophoresis was performed at 110 V for 20–50 minutes. The resulting gel images were captured using a blue transilluminator, Azure C150 (Azure Biosystems).

##### 2.2.2. Size Measurements with Dynamic Light Scattering (DLS)

Hydrodynamic diameters of the three ICG-based probes were measured via DLS using a Malvern Zetasizer Nano ZS instrument. 5-arm DNA nanostructure was folded at 1 μM and DLS measurement was performed without further dilution. The concentration of 50 μM was chosen for JAAZ particles and ICG NBs for the final data collection. A volume of 100 μL of each sample was loaded into disposable plastic cuvettes and size measurements were taken with 173° backscatter.

##### 2.2.3. Scanning Electron Microscopy (SEM) evaluation

SEM was used to confirm size measurements and assess the morphology of JAAZ and ICG-NBs. To prepare the ICG-NB samples, 40 μL of diluted suspension of the samples was placed on a metal slide covered in carbon tape. The slide was kept in an inverted position and freeze-dried with at least 10 minutes of liquid nitrogen and 24 hours of lyophilization. Samples were then sputter-coated via a Coxem, SPT-20 device using gold for 100 seconds at 3 mA (70). SEM was performed using a Low-Vacuum Scanning Electron Microscope (JSM-IT500HR LV SEM, JEOL) at magnification of x60,000, imaged with 15 kV of accelerating voltage.

JAAZ particles were deposited and vacuum dried on silicon wafers and sputter-coated with gold using a Denton Desk V instrument for 30 seconds at 2 mA and imaged with 1-3 kV of accelerating voltage. The SEM imaging was performed using a JEOL JSM-7200F instrument.

##### 2.2.4. Optical Characteristics Measurements

A UV-Visible-NIR plate reader was utilized for the absorbance spectra measurements of all three newly developed ICG-based contrast platforms (Tecan Safire 2 Multi-Detection Plate Reader) for the wavelength range of 500-900 nm.

The fluorescent measurements were performed for the wavelength regime of ∼775–850 nm via a fluorescence plate reader (Molecular Devices Spectramax Gemini Em Fluorescence Microplate Reader). Moreover, the equivalent ICG concentration of the probes was measured using a standard curve plotted for fluorescence and absorbance readouts of at least 6 free ICG concentrations of the range of 500 nM-5 µM and 25-150 µM respectively.

##### 2.2.5. Photoacoustic Signal Acquisition

###### 2.2.5.1. Custom Photoacoustic Setup

The samples were loaded into PTFE tubing with an internal diameter of 100 µm and placed in DI water within a transparent acrylic chamber. A 35 MHz focused piezoelectric transducer was placed 12 mm above each sample which corresponds to the transducer’s focal distance. Photoacoustic signal optimization for each sample was achieved by moving a fine x–y stage over the samples, while the transducer and laser positions remained fixed. A tunable pulsed laser irradiated the samples at an angled incidence. The laser system (Phocus Mobile, Opotek) operates within 690 and 950 nm, delivering around 58 mJ/cm² per 5-nanosecond pulse at a frequency of 10 Hz. The signals were read by the transducer, which is subsequently amplified by a 20/40 dB amplifier (HVA-200 M, Femto). The detected and amplified photoacoustic signal is then processed through a lock-in amplifier (Zurich Instruments, Zurich, Switzerland), synchronized with each laser pulse using a photodiode located near the laser source. Further signal averaging and processing was carried out through MATLAB software.

###### 2.2.5.2. Vevo-F2 Preclinical Imaging System

The samples were first inserted into polyurethane (PU) tubing and stabilized in Vevo Contrast Agent Phantom filled with DI water. A 3.5 mm-thick layer of organic chicken breast tissue was positioned between the tubing and a 29 MHz transducer (center frequency: 20 MHz; bandwidth: 15–29 MHz), maintaining an overall distance of approximately 1 cm. The transducer’s Fiber Jacket cavity was filled with degassed clear ultrasound gel, and the acquired PA signal intensity was averaged along the length of the tube (Figure S8). The Vevo® LAZR-X has a tuning range of 680 - 970 nm and it can deliver ≥ 36 mJ of energy over the range. Utilizing Spectro mode, absorption profile of each contrast agent was measured to define the peak absorbance wavelength for the intended data acquisition (795 nm for ICG and 5-arm DNA ICG probes, 790 nm for ICG-nanobubbles, and 875 nm for JAAZ). As for the photobleaching data collection, the signal was recorded over 2970 frames whereas for the SNR calculations the first 100 frames were taken into consideration. Raw data analysis and quantification were performed using Vevo software (Figure S6). Subsequent data fitting was conducted in MATLAB using a single exponential decay model with an offset (Equation 1). SNR was calculated as the mean signal intensity within regions of interest (ROIs) over the frames (first 20 seconds of laser exposure), divided by the noise, defined as the standard deviation (SD) of signal intensity in a separate ROI placed in the surrounding chicken tissue representing the background (Equation 2). A negative control of a tube injected with PBS was used for this assessment.

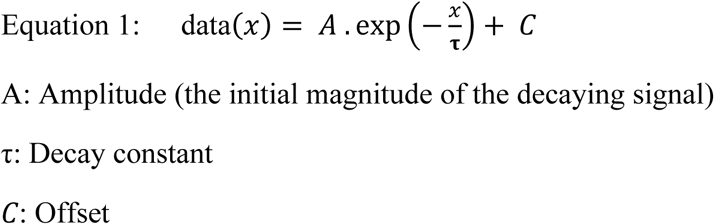

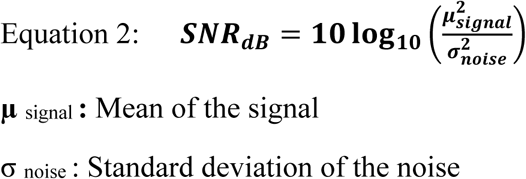

##### 2.2.6. Nanoparticle Tracking Analysis (NTA) with Zetaview® for JAAZ Particle Concentration

The number of JAAZ particles was determined using NTA with the Zetaview®, for sample of JAAZ at an equivalent ICG concentration of 300 μM (Figure S9). Samples have been diluted 20 or 200 times in water and were loaded in the instrument using a 1 ml syringe. Parameters have been adjusted as follows: scatter sensitivity 60, shutter 300 and frame rate 7.5 fps. The particle concentration of the sample was measured at an average of 3.5 × 10^9^ particles/mL which results in a JAAZ particle molar concentration of 5.81 pM after dividing the particle quantities by the Avogadro Number (6.022 × 10^23^). This relation can be applied to JAAZ samples with varying equivalent of ICG concentrations including 60 μM which was used for the probe concentration comparison (Table 2).

**Table 2.**
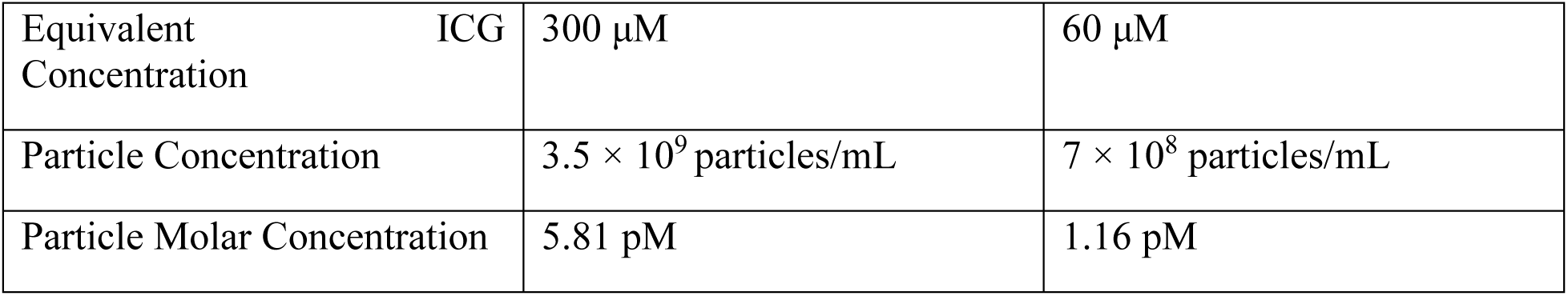
Particle count and molar concentration of JAAZ measured using nanoparticle tracking analysis with the Zetaview® for a reference sample containing 300 μM of equivalent free ICG, with values calculated for the experimental sample (60 μM ICG).

##### 2.2.7. Nanobubble Size, Concentration, and Buoyancy Characterization

The ICG NBs were characterized by size, concentration, and buoyancy using resonant mass measurement (RMM) (Archimedes, Malvern Panalytical Inc. Westborough, MA). NBs were flowed through a nanosensor (0.0001 - 2 mm) that was pre-calibrated with NIST traceable polystyrene beads (diameter: 565 nm). ICG NBs were diluted 1000-fold in 1x PBS (pH 7.4) before collecting the measurements. The molar concentration was calculated similarly to the JAAZ particles. A total of 500 particles were measured for each run (n = 3) (Table 3).

**Table 3.**
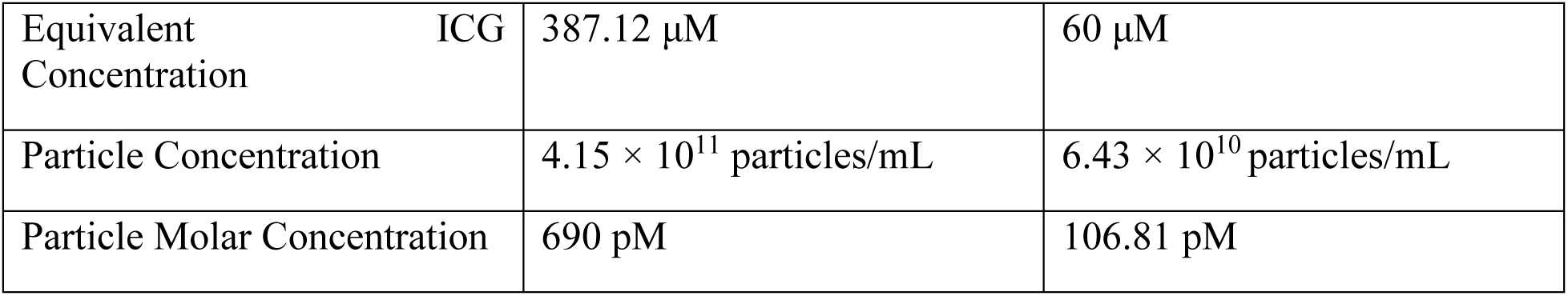
ICG NBs particle and molar concentrations measured with RMM at 387.12 μM ICG concentration, calculated for 60 μM experimental samples.

The bubbles size confirmed with RMM was 239 +/- 2.94 nm which is in agreement with the DLS measurements. The buoyancy of the developed particles was measured at 92.4% (Figure S3).

## Author Contributions

Conceptualization, P.V.C., R.V.; Methodology, M.H., G.G., P.V.C and R.V.; Nanoparticles design and synthesis, M.H., S.S., A.L., D.W., P.N., E.C.A., A.E. and R.V.; Nanoparticles characterization, M.H., S.S., E.R., R.V. All *in vitro* experiments and imaging experiments M.H., G.G.; funding acquisition, P.V.C, and R.V.; M.H. wrote the first draft of the manuscript. All authors contributed to editing the manuscript. All authors have given their approval to the final version of the manuscript.

## Funding Sources

Funding for this research was provided via a National Science Foundation (Award #2128821), a Virginia Innovation Partnership Corporation (VIPC), Commonwealth Commercialization fund grant for Higher education (award #CCF23-0092-HE) and a seed grant from the Office of Research, Innovation, and Economic Impact (ORIEI) at George Mason University (award #G00002563) to P.V.C and R.V.

## Notes

S.S., P.V.C and R.V. have a patent pending on the JAAZ nanoprobes. All other authors declare no competing interest.

## ACKNOWLEDGMENT

We acknowledge the National Science Foundation (NSF) for funding and supporting this study via the award number 2128821, the VIPC for funding and supporting this study via the CCF award number CCF23-0092-HE and the ORIEI at GMU for funding and supporting this study via the award number G00002563. We also acknowledge assistance provided by Dr. Dylan V. Scarton and Bijan Chamanara for their help with the probes’ characterization processes.

## ABBREVIATIONS

NIR, Near infrared; FDA, Food and Drug Administration; DNA, Deoxyribonucleic Acid; ICG, Indocyanine Green; PA, Photoacoustic; PAI, Photoacoustic Imaging; CA, Contrast Agent; 2D, 2 Dimensional; 3D, 3 Dimensional; IJAs, ICG J-aggregates; NP, Nanoparticle; JAAZ, Azide-modified ICG J-aggregates; NBs, Nanobubbles; MBs, Microbubbles; DLS, Dynamic Light Scattering; SEM, Scanning Electron Microscopy, PBS, Phosphate Saline Buffer; SNR, Signal to Noise Ratio; PTFE, Polytetrafluoroethylene; PU, Polyurethane; sO₂, Oxygen Saturation; CEUS, Contrast-enhanced Ultrasound Imaging; PFCs, Perfluorocarbons; FBS, Fetal Bovine Serum; DX, Double-crossover; DIVIN, DNA Integrated Voltage Indicating Nanoparticle; AGE, Agarose Gel Electrophoresis; 1 kb+, 1 Kilobase Plus; bp, Base Pair; RMM, Resonant Mass Measurement; UV, Ultraviolet; IV, Intravenous; DI, Deionized; ROI, Region of Interest; ssDNA, Single Strand DNA; PDI, Polydispersity Index; SD, Standard Deviation; NTA, Nanoparticle Tracking Analysis; NIST, National Institute of Standards and Technology; NBCI, National Center for Biotechnology Information; BLAST, Basic Local Alignment Search Tool; NSF, National Science Foundation; VIPC, Virginia Innovation Partnership Corporation; CCF, Commonwealth Commercialization Fund; ORIEI, Office of Research, Innovation, and Economic Impact;

