## Supplementary Information for "Synthesis and Characterization of ICG-based Near-infrared Photoacoustic Contrast Agents"

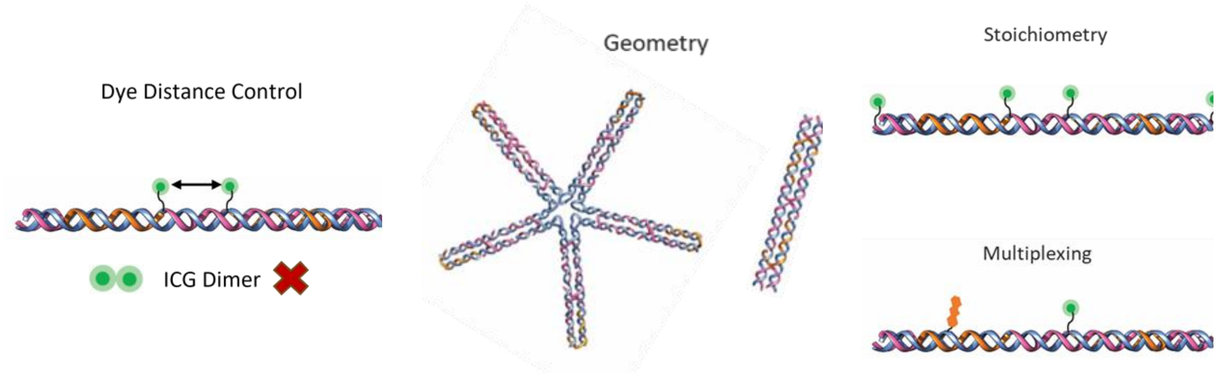


**Figure S1**. DNA-ICG Platform: Precise nanoscale organization enables controlled placement of ICG and targeting moieties, minimizing dye aggregation and supporting modular, cell-specific labeling across a wide range of scaffold shapes and sizes.

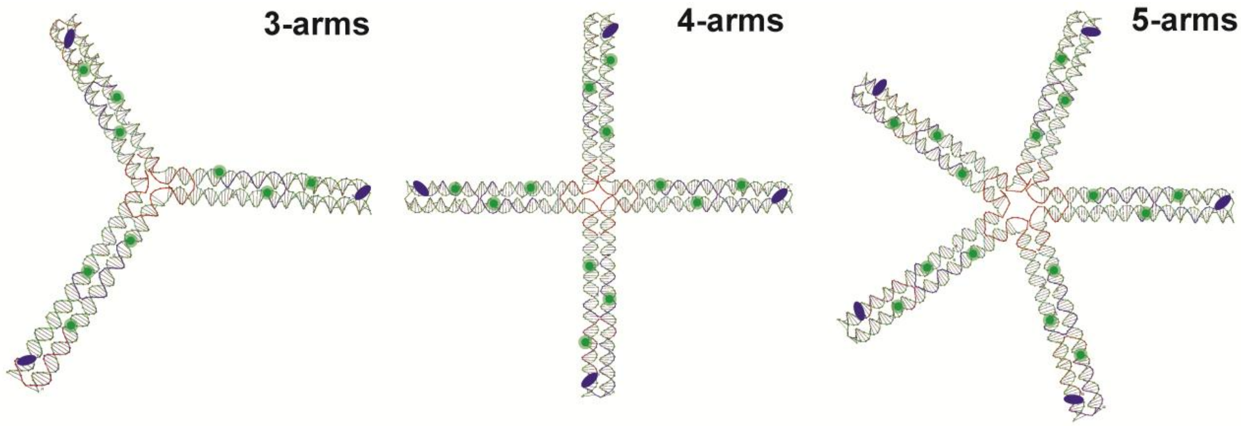


**Figure S2.** 2D DNA nanoparticles design: 3,4 and 5-arm 2-dimentional DNA ICG nanostructures bear 9, 12 and 15 ICG molecules respectively (green circles) while maintaining minimum inter–dye distance to prevent aggregation. The 3-arm model offers 3 sites for the conjugation of targeting moieties while 4-arm and 5-arm can support 4 and 5 ligands on their structures (navy blue ovals).

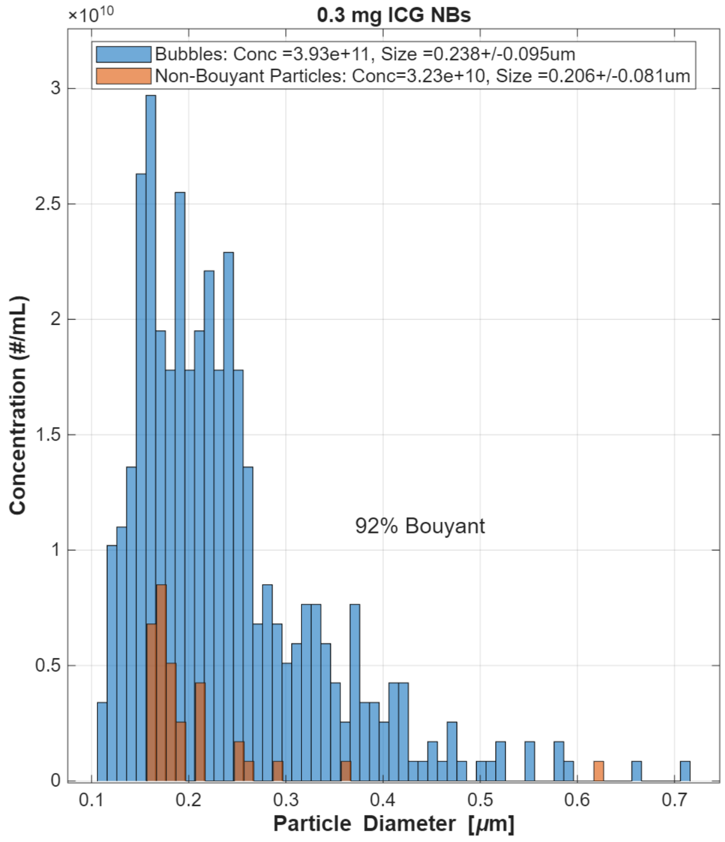


**Figure S3:** Resonant mass measurement sample result for ICG NBs: A size distribution, bubble concentration and buoyancy analytical outcome for ICG nanobubbles is depicted in the figure above.

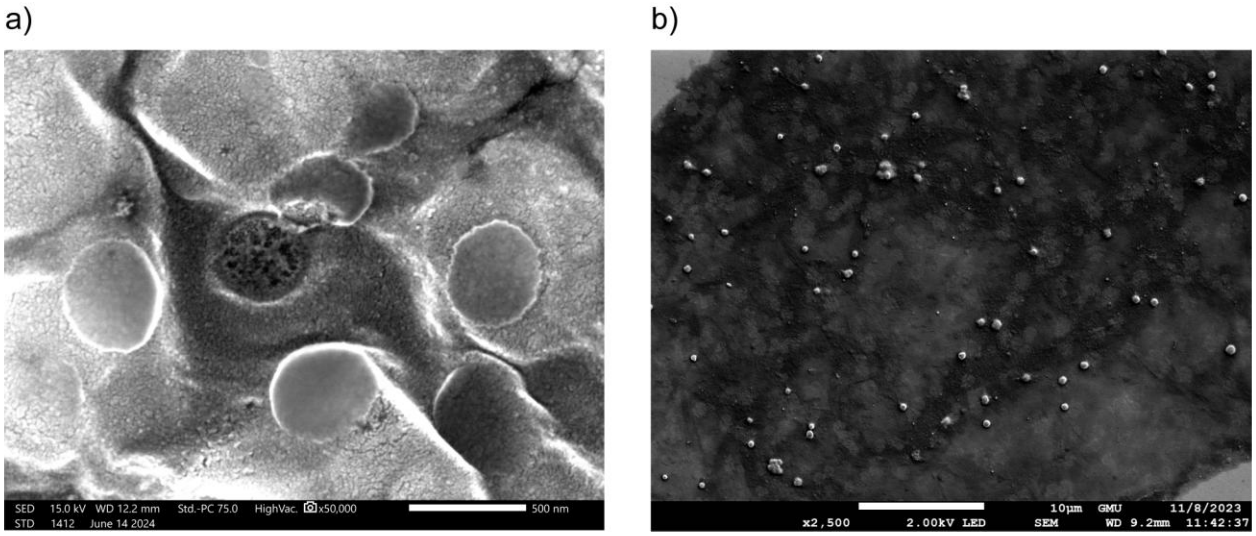


**Figure S4.** SEM images of ICG nanobubbles and JAAZ: a) An SEM image of the lipid nanobubbles including the one in the middle that was disrupted during scanning electron microscopy, due to damage induced by the high accelerating voltage. Scale bar: 500 nm. b) The overall morphology of JAAZ particles can be observed in the low-magnification image, providing a broader view of the sample. Scale bar: 10 µm.

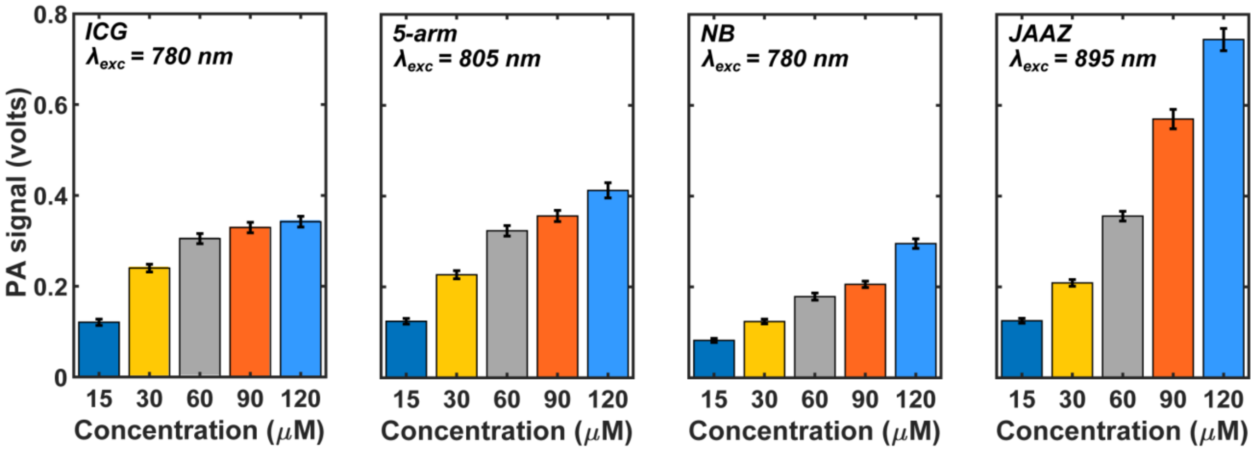


**Figure S5:** Photoacoustic Signal Amplitude Measurements in PBS: Similar to the data collected for the samples in blood, JAAZ produced the strongest signal at high concentrations, depicting more than a twofold increase at 120 µM compared to ICG. DNA-ICG demonstrated slight improvement in signal amplitude over ICG with visible increasing trend when the concentration rises which cannot be observed for free ICG. ICG-nanobubbles in PBS, however, showed weakened signal at all concentrations in comparison with ICG, although a concentration-dependent signal enhancement can be seen. As stated before, this contrast between ICG-NBs performance in blood and PBS suggests that the acoustic response improvement for NBs in blood is possibly attributed to direct interactions or association with red blood cells via hitchhiking effect.

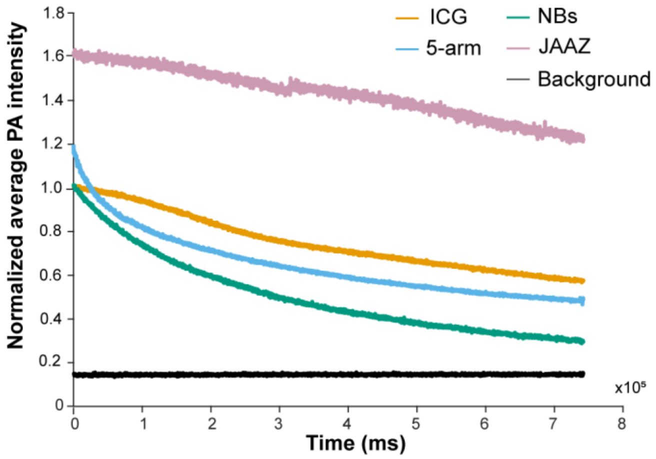


**Figure S6:** Raw PA signal magnitude data for the developed ICG-based nanostructures over approximately 12 minutes, normalized to the initial PA signal intensity of ICG at time zero ($t_0$), without any model fitting. All agents were prepared in PBS at an equivalent ICG concentration of 50 µM. The black trend line denotes the background signal.

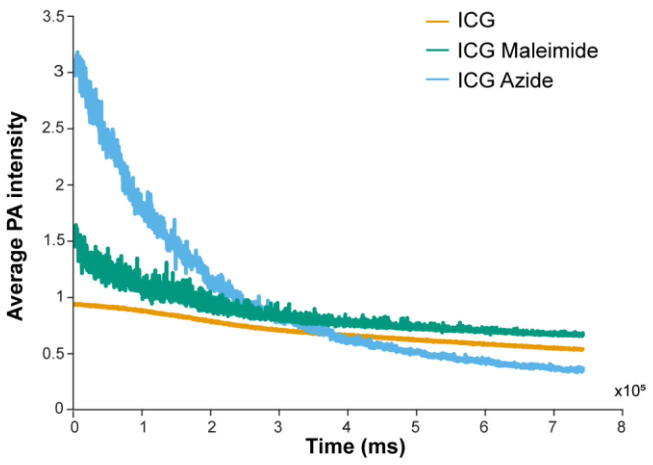


**Figure S7.** ICG, ICG Azide and ICG Maleimide PA signal intensity over time: ICG Azide is used to produce JAAZ particles (in addition to ICG) and ICG Maleimide is the molecule exploited for templating ICG on our developed DNA nanostructures. As illustrated in the figure, adding functional groups to ICG affects signal amplitude, decay rate, and fluctuations. Both ICG Azide and Maleimide initially show elevated signal intensity while exhibiting a rapid signal decay over time compared to ICG without functional groups. This outcome indicates that for 5-arm DNA-ICG structure, the initial amplified signal and its decay may be attributed to the use of ICG Maleimide in its synthesis process as well. Moreover, ICG Azide may have assisted JAAZ particles in enhanced signal amplitude in the beginning, but their presence adversely affects signal decay pace over time. Nevertheless, JAAZ manages to demonstrate superior photostability which stems from its aggregated structure.

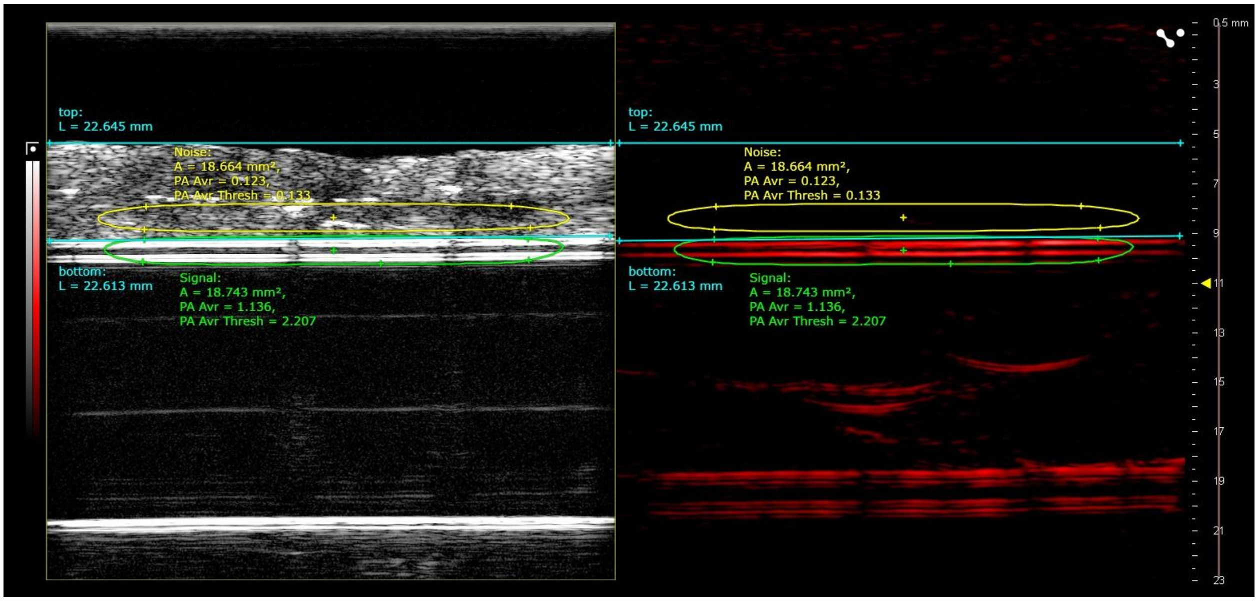


**Figure S8:** Ultrasound and photoacoustic images of the sample setup acquired using Vevo software: The region outlined in green indicates the sample inside the tubing (signal area), while the yellow-bordered area represents the noise region, positioned over the chicken tissue. The organic chicken breast tissue has a 3.5 mm thickness whereas the overall distance between the sample and the transducer is 1 cm.

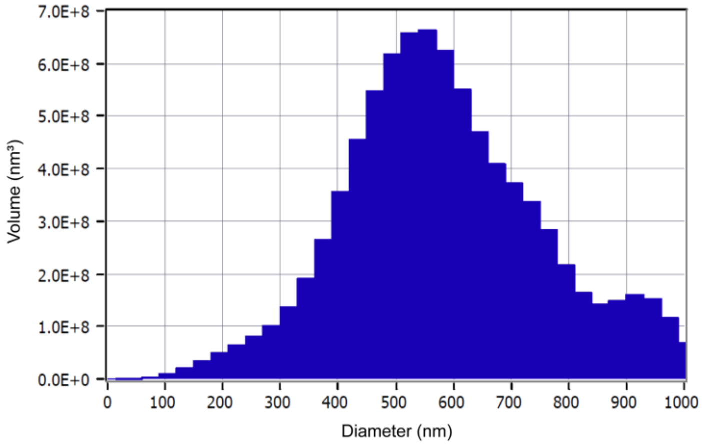


**Figure S9:** Zetaview measurement sample results for JAAZ: A size distribution outcome for Azide-modified ICG J-aggregates is depicted in the figure above.

**Table S1.** Non-modified DNA oligonucleotides used to assemble the 5-arm probe.

| Strand | Ratio for Folding | Sequence | Length (nts) |
|---|---|---|---|
| Core - 1 | 1/37 | TCC CTG CAC TTT TTC TAA GGA AGC TAC CAA TAT TTT GTT TCC GAG TCT CAC GTC TGA CCT TTT TGG TAG ATT GCC | 75 |
| Core - 2 | 1/37 | ATG CAT AGA GTT TTC GAG TCA GCG AAA AGT ATG AGT TTT TTT TAA GTA TG | 50 |
| Arm - 1 | 5/37 | TAG TAG GAC GAA GGC TCG TAT CCG TCC AAC TA | 32 |
| Arm - 2 | 5/37 | AGA CCA GCG TAC GTC TAC CAG TTT TTT TAA GG | 32 |
| Arm - 3 | 5/37 | TTC GTC CTA CTA CCT TAA AAT GCG GTG CGG TTG CGT GTG GGT TGC AAA GAC | 51 |
| Arm - 4 | 5/37 | CTC GTT ATC CTA ATT CAT ACA TGG TCT TTG C | 31 |
| Arm - 5 | 5/37 | AAC CCA CAC GCA ACC GCA CCG CAT ATG TAT C | 31 |
| Arm - 6 | 5/37 | CAT GTA TGA ATT AGG ATA ACG AGG ATA CAT AAA ACT GGT AGA CGT AC | 47 |
| Core - 3 | 1/37 | GCT GGT CTT AGT TGG AGG CAA TCT ACC AGG TCA GAC GCG GAT ACG AGC C | 49 |
| Core - 4 | 1/37 | CTG GTC TTA GTT GGA TCG CTG ACT CGC TCT ATG CAT CGG ATA CGA GCC | 48 |
| Core - 5 | 1/37 | GCT GGT CTT AGT TGG ACA TAC TTA AAA CTC ATA CTT TCG GAT ACG AGC C | 49 |
| Core - 6 | 1/37 | GCT GGT CTT AGT TGG AAG CTT CCT TAG AGT GCA GGG ACG GAT ACG AGC C | 49 |
| Core - 7 | 1/37 | GCT GGT CTT AGT TGG ATG AGA CTC GGA AAA TAT TGG TCG GAT ACG AGC C | 49 |

**Table S2.** Modified DNA oligonucleotides used to assemble the 5-arm probe.

| Strand | Ratio for Folding | Sequence | Length (nts) |
|---|---|---|---|
| Arm - 1 | 5/37 | TAG TAG GAC GAA GGC TCG TAT CCG TCC AAC TA-ICG | 32 |
| Arm - 4 | 5/37 | CTC GTT ATC CTA ATT CAT ACA TGG TCT TTG C-ICG | 31 |
| Arm - 6 | 5/37 | CAT GTA TGA ATT AGG ATA ACG AGG ATA CAT AAA ACT GGT AGA CGT AC-ICG | 47 |
